# Optimisation of functional network resources when learning behavioural strategies for performing complex tasks

**DOI:** 10.1101/2020.06.17.156570

**Authors:** Richard E. Daws, Gregory Scott, Eyal Soreq, Robert Leech, Peter J. Hellyer, Adam Hampshire

**Author notes:** **Corresponding author:** Name: Richard E. Daws.

## Abstract

We developed two novel self-ordered switching (SOS) fMRI paradigms to investigate how human behaviour and underlying network resources are optimised when learning to perform complex tasks with multiple goals. SOS was performed with detailed feedback and minimal pretraining (study 1) or with minimal feedback and substantial pretraining (study 2). In study 1, multiple-demand (MD) system activation became less responsive to routine trial demands but more responsive to the executive switching events with practice. Default Mode Network (DMN) activation showed the opposite relationship. Concomitantly, reaction time learning curves correlated with increased connectivity between functional brain networks and subcortical regions. This ‘fine-tuning’ of network resources correlated with progressively more routine and lower complexity behavioural structure. Furthermore, overall task performance was superior for people who applied structured behavioural routines with low algorithmic complexity. These behavioural and network signatures of learning were less evident in study 2, where task structure was established prior to entering the scanner. Together, these studies demonstrate how detailed feedback monitoring enables network resources to be progressively redeployed in order to efficiently manage concurrent demands.

**Highlights:**

- We examine the optimisation of behaviour and brain-network resources during a novel “self-ordered switching” (SOS) paradigm.
- Task performance depended on generating behavioural routines with low algorithmic complexity (i.e., structured behaviours).
- Behaviour became more structured and reaction time decreased as SOS was practised.
- As behaviour became more structured, activation in multiple-demand regions decreased for simple trial events but increased for executive switching events
- Increases in between-network functional connectivity correlate with reaction time decreases.

## Introduction

The ability to rapidly learn complex tasks is a definitive feature of human cognition. Studies of cognitive control suggest that the human brain achieves this by rapidly adapting to encode behavioural routines (Finc et al., 2020; Khambhati et al., 2018), also known as a ‘task set’, comprising the combinations of rules, strategies and stimulus-response mappings that enable optimal performance (Sakai, 2008). This encoding is typically associated with distributed activation of the multiple-demand (MD) system (Duncan, 2013).

During instruction based learning (IBL), generating and establishing a simple routine imposes a cognitive demand that rapidly decreases with practise. This learning can be accompanied by a decay in MD activation, particularly in anterior prefrontal cortex (aPFC), as reaction time (RT) decreases (Hampshire et al., 2016, 2019). The established behavioural routine can be robustly identified by task-specific multivariate patterns of the brain’s functional connectivity (FC) (Soreq et al., 2019, 2021) and these patterns are predictive of behavioural performance (Rosenberg et al., 2016; Soreq et al., 2021). As individuals learn a task, FC is dynamic and becomes more stable with increased FC within cortical networks (Hampshire et al., 2016) and in their connectivity with subcortical regions (Antzoulatos & Miller, 2014; Bassett et al., 2011).

In everyday life, the optimal behavioural routine is often not evident when first performing tasks. This can be due to a lack of clear instruction, the inherent complexity of the task, or shifting demands. Consequently, it is necessary to monitor outcomes and progressively update behavioural routines in order to optimise behaviour (Sharp et al., 2010; Swick & Turken, 2002). This updating of routines is critical for flexible behaviour (Ionescu, 2012); however, it comes with a cost and is associated with increases in reaction time (RT) and heightened MD activation (Daws et al., 2020; Hampshire & Owen, 2006). Furthermore, it often is necessary to juggle multiple concurrent tasks, or to manage sub-routines of complex tasks, and in the process, to monitor multiple outcomes. Increased task complexity poses a challenge for flexible cognitive systems due to their intrinsic limitations in capacity (Duncan, 2013). Under such conditions, humans have a tendency to impose hierarchical structure to their behavioural routines (Fallon et al., 2013).

This tendency towards self-ordered structure is underpinned by decisions regarding when and in what order to switch attention between competing tasks. Consequently, switching paradigms provide a natural choice for studying the cognitive control processes that enable flexible human behaviour. However, task-switching paradigms typically control when switches occur (Kiesel et al., 2010), which does not capture the self-ordered nature of real-world human behaviour (Hampshire & Owen, 2006). Despite this limitation, there is evidence that cognitive demand is minimised with learning and by structuring behaviour, where possible. Task-switching imposes a considerable cognitive demand but switching-costs to RT will decrease with practise (Yeung & Monsell, 2003), although a residual cost persists in humans even after tens of thousands of trials (Stoet & Snyder, 2007). Moreover, switching-costs decrease with preparation time and also when switching events are predictable (Rogers & Monsell, 1995). During voluntary task-switching (VTS), participants are less likely to switch under high cognitive load (Demanet et al., 2010) and can exhibit a repetition bias despite being instructed to evenly and randomly switch between tasks (Arrington & Logan, 2004, 2005).

These findings indicate that both learning and the structuring of behaviour are important mechanisms for minimising cognitive demand when managing competing task demands. Here, we investigate this further using functional MRI (fMRI) and two novel “self-ordered switching” (SOS) paradigms. The SOS paradigm requires participants to learn through a process of feedback-driven trial and error how to manage their time between two tasks, each of which has two rules that the participants select. To score maximum points, the participant must spend time on all four sub-tasks. In two studies, the concurrent optimisation of brain function and behaviour was examined as participants either actively learnt SOS demands based on multiple feedback elements (study 1) or, as a control condition, performed SOS with minimal feedback only and after receiving extensive pre-training (study 2).

We tested the hypothesis that in study 1, where participants were performing the task for the first time and with detailed feedback, self-ordered behaviour would optimise with practise. It was predicted that RT would decrease and that behavioural routines would become increasingly structured. Using fMRI measures of network activation and FC, we hypothesised that brain function would vary as behaviour optimised. Specifically, we predicted that MD activation would decrease with practise for the simple trial events, but that these learning curves would be less rapid for the more demanding switching events. We predicted that FC increases between functional networks associated with cognition would accompany the learning of optimal behavioural routines, reflecting more efficient network processing. Finally, we expected that in study 2, where the task had been pre-trained and feedback was minimal, that participants would show little or no behavioural learning effects.

## Methods

### Participants

31 healthy right-handed participants were recruited in total: **Study 1** - n=16, mean age 24.75 years, range 19-40, 10 female; **Study 2** - n=15 mean age 26.47 years, range 19-42, 11 female. Approval for this study was received by the Cambridge university research ethics committee and participants gave written informed consent. Participants were reimbursed £20 for their time.

### Self-ordered switch (SOS) paradigms

We designed two SOS paradigms that required participants to manage their time between two concurrent tasks that each had two subordinate rules in order to gain maximum points within 20 minutes. There was a matching task (MT) and an odd one out (OOO) task, both involving simple visual discriminations of objects (Figure 1a). The subordinate rules of each task were differentiated by the requirement to apply the rules to either colour or shape. During each trial, arrays of simple colour-shape objects were presented, and participants made a visual discrimination response based on the current task and rule.

**Figure 1.**
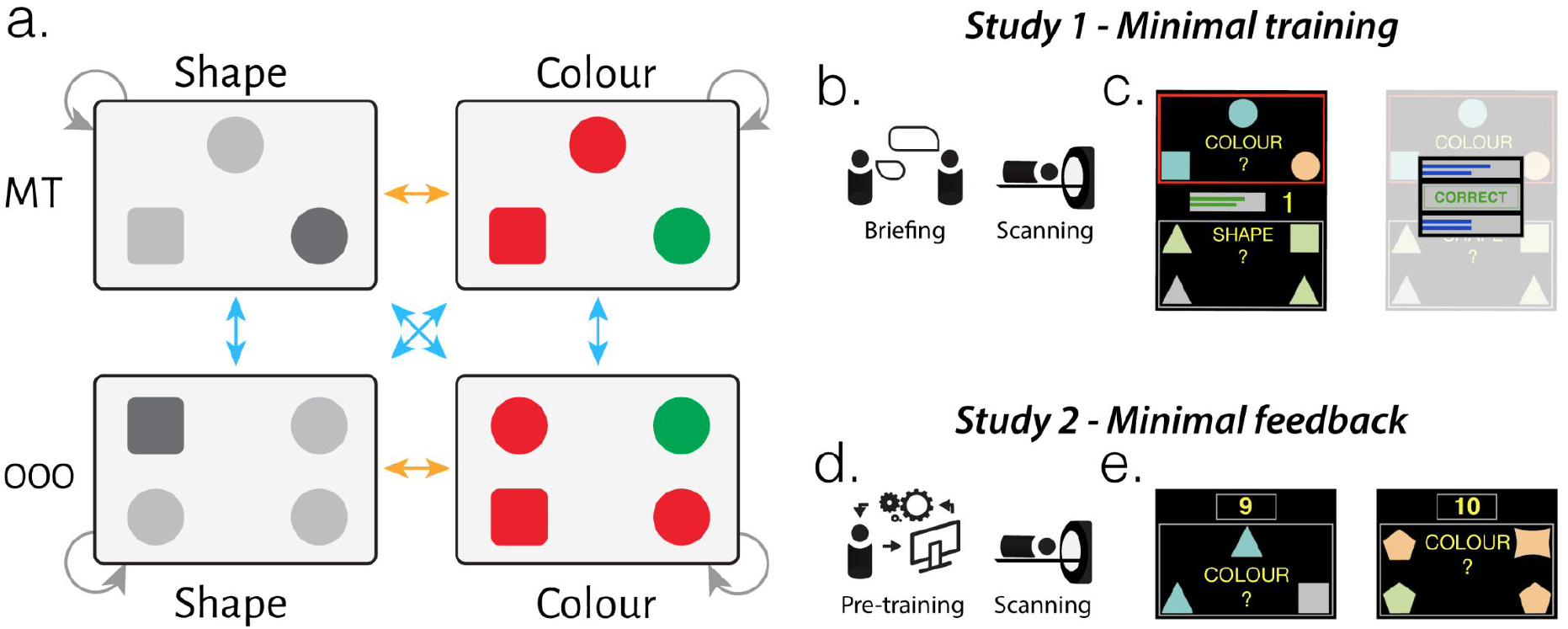
Self-ordered switching (SOS) experiments. a) Schematic of the two tasks (MT - Matching task, OOO - Odd one out) and the two rules (colour, shape) of the SOS paradigm. Grey arrows represent repeat-trials where the same condition was repeated. Task-switching and rule-switching are represented by the blue and orange arrows, respectively. b) In study 1, SOS was performed after minimal training with a short demo. c) Both tasks were simultaneously displayed on the screen and hierarchical response feedback was presented with progress meters on each task overall as well as each of the task’s subordinate rules. d) In study 2, SOS was performed after extensive training with full hierarchical feedback and then performed in the scanner with minimal feedback involving the overall score only (e).

Critically, in order to keep gaining points, participants had to organise their time across all four sub-tasks. This meant that they chose whether to repeat the same condition (repeat-trial) or to switch between the tasks or the rules (task-switch, rule-switch). Responses were made with a button positioned under the right hand. Each trial continued until a response was made using either a left or right button press. The task and rule task did not change between trials unless the participant opted to task-switch or rule-switch by pressing the up or down button respectively.

When performing MT trials, a target and two probe objects (each with a shape and colour) were presented, and the participant indicated via a left or right button press which of the two probes matched the target given the currently selected shape or colour rule. Similarly, during OOO trials, a 2×2 object grid was presented and the participant indicated with a left or right button press which side of the display contained the object that differed from the others given the currently selected colour or shape rule.

The task and rule discrimination criteria were designed to be simple enough that individual trials could be performed with high accuracy and minimal effort. This ensured that individuals could focus on the executive requirement to learn how to divide their time across the tasks and rules in order to gain maximum points (see detailed criteria below). In both studies, SOS was continuously performed inside the scanner for 20 minutes with no resting periods. The studies differed with respect to the training administered and the response feedback received during the scanning session.

In study 1, participants played a short demo, during which they were required to undertake each of the four task-rule combinations and each of the switch types once (rule or task switch). On screen instructions were sequenced to orient the participant towards the feedback elements (described below) and controls for responding and switching. They then entered the scanner and undertook the SOS paradigm in a single continuous 20 minute acquisition. Both tasks were displayed on the screen with the MT at the top and the OOO task at the bottom (Figure 1c). The currently active task was indicated with a red border, the inactive task had a grey border (these border colours flipped when the task-switch button was pressed). The active rule was displayed with a text cue in the centre of the task panel (“COLOUR” or “SHAPE”, in yellow) and flipped when the rule-switch button was pressed. On each trial, new stimuli were displayed, and participants were able to repeat the condition or could task-switch and/or rule-switch once prior to performing the trial within the selected panel. A feedback array was then immediately displayed for 1.5 seconds followed by the stimuli for the next trial (Figure 1c).

Participants learnt to optimally sequence their responses based on a hierarchically arranged set of performance meters (supplemental Figure 1a-b). These meters were as follows. (1) An overall score meter was located at the centre of the screen alongside (2) two meters indicating the number of points currently accumulated for each of the two tasks. These were presented continuously on the screen. Additionally, the feedback array after each trial comprised (3) text indicating positive or negative feedback for the present trial response (“CORRECT” in green or “INCORRECT” in red) and (4) four meters, showing points currently accumulated on each of the sub-task (i.e., colour and shape for MT vs OOO).

These score elements interacted hierarchically. Specifically, sub-task meters would increase after a correct response to the corresponding condition (e.g., a correct response to a MT colour trial) and would decrease if an incorrect response was made. All meters ranged between one to ten and were initialised with five points at the start of the experiment.

Within each MT and OOO task, the aim was to respond to, and switch between, the task’s colour and shape rules in a sequence that would build up both rule meters to the maximum level of ten (supplemental Figure 1c-e), at which point, the two subtask meters would reset to five, and the corresponding high-order task meter and overall score incremented by 1. To encourage switching, if one subtask meter got further than two points ahead of the other, then the other meter decreased by a point (supplemental Figure 1d).

Switching between tasks was encouraged in a similar way to rule switching as the inactive task meter would decrease if the active task meter was two or more points ahead. Successfully dividing time between each task would build up both task meters, thereby increasing the overall score. When both task meters reached the maximum level of ten, both meters would reset to the initial position and a +10 point bonus was awarded to the overall score (supplemental Figure 1f).

Given that switching would be costly in terms of both effort and time, and the aim was to score as many points as possible per unit time, optimal behaviour could be defined as having a routine that used the minimal number of switches required to ensure sufficient dividing of time between each task’s sub-rules and between each task. Those who learnt the most efficient way of doing this, via trial and error, would be expected to score the most points during the experiment. An example sequence of events that a participant could perform is described in the supplementary materials.

In study 2, participants performed SOS with the same instructions, stimuli, rules and tasks; however, there were modifications in the presentation of the tasks, training and feedback. Most critically, immediately prior to the scanning session, participants received a 20 minute training session as opposed to a minimal demo. This extensive training session was designed to enable participants to arrive at a structured routine for performing the task prior to the scanner session. Additionally, the task and sub-task feedback scales from study 1 were only present during the training. Therefore, the only feedback available in the scanner was represented by the overall score number. There also were no bonus points awarded on increasing both task scales to their limit whilst in the scanner. These modifications were designed to minimise the information that the participant could use to further optimise their behavioural routine inside of the scanner whilst ensuring that they had sufficient information to remain engaged. Finally, as opposed to presenting both tasks simultaneously (study 1), only the currently selected MT or OOO task was presented on the screen (Figure 1e). This modification was intended to further control for activity related to switching between spatial locations as opposed to tasks.

### Behaviour

The RTs from correct trials were separated into three categories: Repeat-trials, rule-switch and task-switch. Repeat-trial RT was the time between the stimuli presentation and the L/R response being made. Rule-switch or task-switch RT was the time between the corresponding switch button being pressed and the subsequent L/R response. For each participant, RTs for each category were summarised using the median from correct trials. The number of trials completed would naturally vary across participants and response accuracy was summarised for each RT category as the percentage of correct responses made. Separate one-way repeated-measures analysis of variance (RM-ANOVA) models were independently conducted for the RT and accuracy measures

Trial conditions could be repeated or switched between as each participant saw fit (Figure 1a). To quantify the extent to which participants structured their behaviour, we estimated the algorithmic complexity (Zenil et al., 2018) of each behavioural timecourse. Algorithmic complexity estimates the size of the smallest algorithm that could generate a given sequence and tends to be smaller for sequences with consistent structures vs. random sequences.

Here, we use the framework, outlined by Zenil et al. (2018), which combines a block decomposition method (BDM) and the coding theorem method (CTM) (Lempel & Ziv, 1976) to provide a reasonable estimate of Kolmorogov-Chaitin complexity (K) (Chaitin, 1966; Kolmogorov, 1968) for large strings in a computationally tractable way. The advantage of BDM is that it provides an estimate of K that not only considers statistical regularities, (e.g., as is captured by Shannon entropy), but it is also sensitive to segments of an algorithmic nature.

Timecourse length naturally varied across individuals, and to account for this, we normalised an individual’s algorithmic complexity by the mean algorithmic complexity calculated from 100 randomly shuffled versions of the individual’s data (qualitatively similar results were obtained without normalisation). NB:-As expected, the shuffled data exhibited significantly greater algorithmic complexity (paired t-test: **Study 1** - t_15_=−9.775, p<0.001, *d*=3.54; **Study 2** - t_14_=−35.329, p<0.001, *d*=14.29).

A more focused behavioural analysis was conducted to examine how RT and algorithmic complexity varied over the course of the 20 minute experiment. Here, the behavioural timecourses were split into three non-overlapping time-windows (6.67 minutes of data), or “Learning stages”. The median RT for each category (correct trials) as well as the normalised algorithmic complexity (using the same procedure outlined above) were extracted from each learning stage.

### fMRI acquisition and preprocessing

Whole-brain blood oxygenation level dependent (BOLD) fMRI recordings were acquired using the same Siemens 3T Trim Trio scanner and following parameters in both studies: TR=2s, TE=30ms, FA=78°, 3mm^3^ voxels, 3.75mm slice-gap, 32 axial slices. 630 volumes were acquired during the 20-minute SOS task run. A T1-weighted magnetization prepared rapid acquisition gradient echo (MPRAGE) sequence was also acquired using the following parameters: TR=2250ms, TE=2.99ms, IT=900ms, FA=9°, 1mm^3^ voxels, 256 axial slices.

fMRI pre-processing was conducted using standard parameters with the Statistical Parametric Mapping 12 (SPM)^1^ toolbox in Matlab 2018b^2^. The first 10 volumes were discarded to account for T1-equilibrium effects. The remaining images were slice-time and motion-corrected, and the mean echo planar image (EPI) co-registered to the tissue segmented T1 image, normalised to 2mm^3^ MNI space and spatially smoothed (full half width maximum = 8 mm kernel).

### fMRI activation modelling

Participant’s voxelwise fMRI data was modelled using General Linear Models (GLM) with the psychological events defined by responses to one of the four trial conditions (MT colour, MT shape, OOO colour, OOO shape) and when one of the two task-switches (MT to OOO, OOO to MT) or four rule-switches (MT colour to MT shape, MT shape to colour, OOO colour to shape, OOO shape to colour) occurred. Errors were infrequent but were captured in an additional predictor. Models in study 1 included an additional set of feedback events for when the rule or task bars de/incremented and when bonus points were awarded. These psychological predictors were convolved with the canonical haemodynamic response function (HRF) with the durations set to 0s.

Head motion and nuisance variables were modelled using the rigid-body realignment parameters, the mean timecourses from the white matter and cerebrospinal fluid tissue segmentations, alongwith their 1st order temporal derivatives. This resulted in 16 parameters. Additional spike regressors were included where framewise displacement (Power et al., 2012) exceeded the voxel acquisition size (3mm).

The 1st-level beta maps were estimated for each psychological predictor reactive to the model intercept (as there were no resting periods). Modelling against this “implicit baseline” results in significant activation differences being relative to the average activation across time.

Additional 1st-level models were created to examine how activation varied over time. Specifically, the psychological predictors were segmented by the three learning stages defined in the behavioural analysis. The trial response events were collapsed into a single predictor, and switching events were collapsed into a single task-switch and a single rule-switch predictor. The same nuisance predictors were used in these models.

The group-level modelling of activation associated with switching involved a voxelwise one-way RM-ANOVA of the beta maps representing the two task-switch and four rule-switch conditions. To examine how switching activation varied over time, switch>repeat contrasts were generated from each learning stage and modelled with a group-level two-way RM-ANOVA with learning stage (3 levels) as the within-subject factor and study (2 levels) as a covariate. All group-level activation maps were initially thresholded at an uncorrected p<0.01, followed by a p<0.05 false discovery rate (FDR) cluster correction.

### fMRI connectivity modelling

Generalised psychophysiological interaction (gPPI) models (Friston et al., 1997; McLaren et al., 2012) were used to examine how connectivity varied as SOS was performed. Specifically, gPPIs were fitted between pairs of mean timecourses from 274 ROIs defined by external atlases that parcelated the cortex, subcortex and cerebellum (Buckner et al., 2011; Fan et al., 2016; Schaefer et al., 2018). gPPI’s were simultaneously estimated for the psychological events in each learning stage and pair of ROIs using the following:

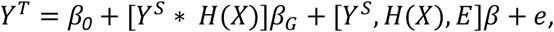

where, *X*, is the matrix of psychological timecourses and, *H(X*), its HRF convolution. *Y^T^* is the target ROI timecourse and, *Y^S^*, the seed ROI timecourse. *E* is the matrix of motion & tissue signal timecourses. *β_G_* are the gPPI betas weights, *β*, the beta weights of no-interest, *β*_0_, is the model intercept and, *e*, the residual. The resulting betas from each pair of ROIs were averaged ([model A->B + model B->A] / 2) and z-scored across learning stage. Subsequently, each dFC estimate represented the change in FC relative to that connection’s baseline FC.

To estimate how gPPIs varied over the learning stages, generalised linear mixed effect (glme) models were fitted to each connection. This mass-univariate approach modelled each connection with the three learning stages as a fixed effect and participants as a random effect. The resulting model set was corrected for multiple-comparisons at a p<0.05 FDR threshold.

## Results

### SOS costs to reaction time decrease with practise

As expected, trials were performed with near-ceiling accuracy (**Study 1** - mean=94.87%, SD=4.18; **Study 2** - mean=94.69%, SD=4.04). Response accuracy did not vary across the repeat-trials, rule-switch or task-switch trials (one-way rm-anova: **Study 1** - F_2,30_=1.218, p=0.310, 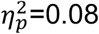; **Study 2** - F_2,28_=1.867, p=0.173, 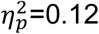).

In contrast, RT varied substantially between the repeat-trials, rule-switch and task-switch trials in both studies (one-way rm-anova: **Study 1** - F_2,30_=48.588, p<0.001, 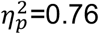; **Study 2** - F_2,28_=52.886, p<0.001, 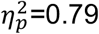). At a finer grain, in study 1 (Figure 2a), task-switching slowed response speed on average by 883ms (SD=538) (paired t-test: t_15_=6.573, p<0.001, confidence interval 95% (CI)=0.60 to 1.17, Cohen’s d=1.64), and rule-switches were 179ms (SD=273) faster (t_15_=−2.630, p=0.019, CI=0.03 to 0.33, d=0.66), relative to repeat-trials (FDR corrected). In study 2 (Figure 2b), response speed slowed by 267ms (SD=143) after rule-switching (paired t-test: t_14_=7.214, p<0.001, CI=0.19 to 0.35, d=1.86), and by a further 362ms (SD=268) after task-switching (t_14_=5.227, p<0.001, CI=0.21 to 0.51, d=1.35) (FDR corrected).

**Figure 2.**
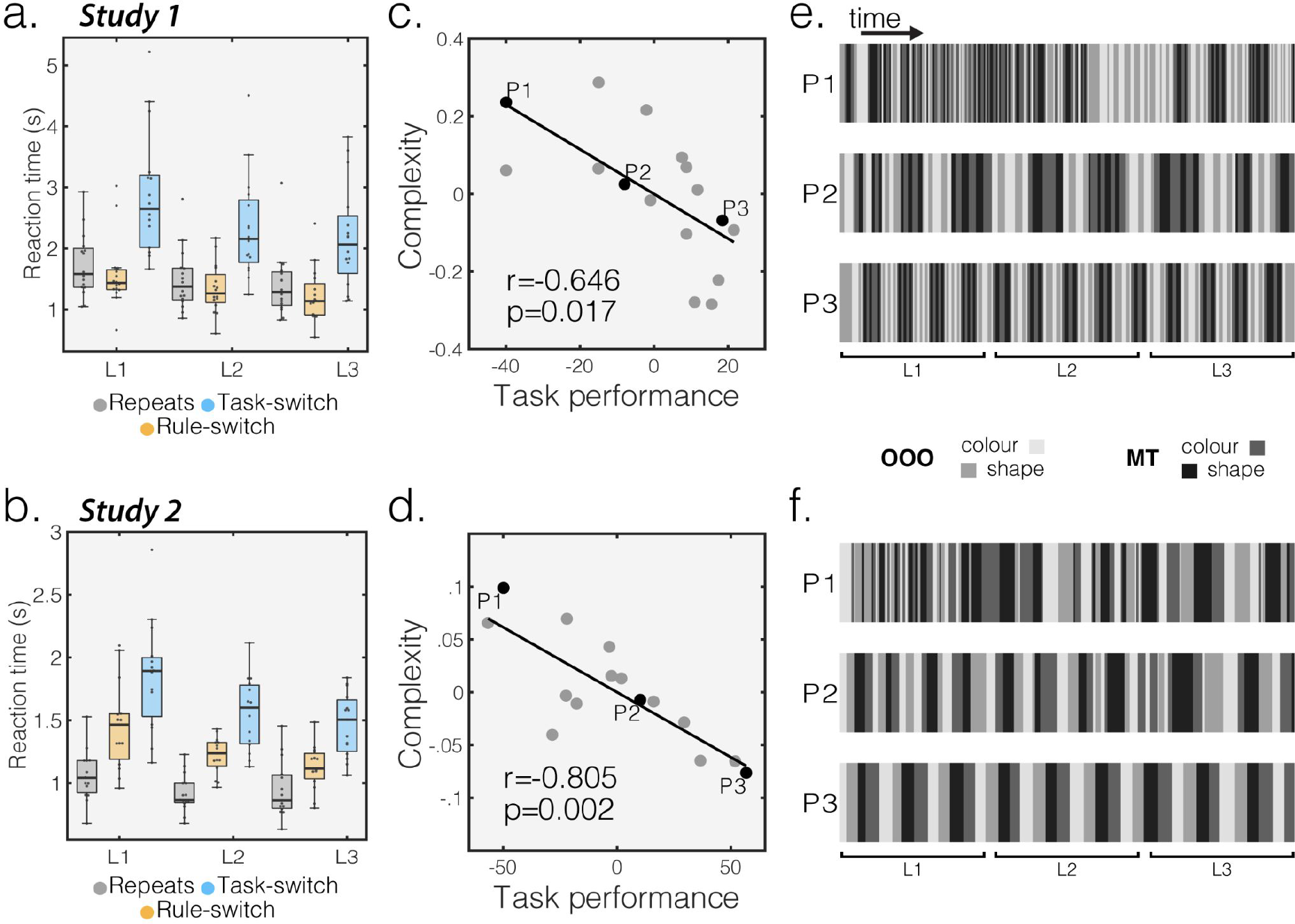
SOS switching costs and behavioural structuring. Switching-costs to reaction time (RT) decreased with practise in study 1 and 2 (a/b). Reaction times (RT) for each condition were summarised within each ~7 minute learning stage (L). Each point represents a participant’s median RT for a condition. Boxplots represent the group median (central line), and the 25-75th percentiles (boxes). c/d) Normalised algorithmic complexity (y-axis) of participants’ SOS behaviour against overall task performance, in terms of points scored (x-axis). The residuals are plotted from a partial correlation controlling for the number of trials completed. For each study, three participants are highlighted and their behavioural timecourses are displayed in e/f. Of those, participant 1 (P1) generated the most complex SOS behaviour and P3 the least.

In order to examine changes in switching-costs with learning, the trials were segmented by the three time-windows, or “learning stages”. Switching-costs for each learning stage were calculated by subtracting the mean repeat-trial RTs from the task-switch or rule-switch trial RTs. Task-switching costs varied over learning stages in both studies (one-way RM-ANOVA: **Study 1** - F_2,30_=3.579, p=0.040, 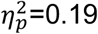; **Study 2** - F_2,28_=5.413, p=0.010, 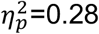), reducing between the 1st and last stage by 369ms (SD=607) in study 1 (paired t-test: t_15_=−2.433, p=0.028, CI=0.05 to 0.69, *d*=0.61), and by 278ms (SD=418) in study 2 (t_14_=−2.570, p=0.022, CI=−0.51 to −0.05, *d*=0.66). Rule-switching costs did not vary significantly across learning stages in study 1 (F_2,30_=0.128, p=0.881, 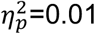), but did in study 2 (F_2,28_=5.543, p=0.009, 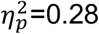) and reduced by 217ms (SD=286) between the 1st and last learning stage (t_14_=−2.945, p=0.011, CI=−0.38 to −0.06, *d*=0.76).

Individual differences in switching costs were examined further by correlating mean switch-trial RTs with the total number of switches performed across participants (supplemental Figure 2a-b). There was a significant negative correlation (Pearson: **Study 1** - r_15_=−0.557, CI=−0.83 to −0.09, p=0.025; **Study 2** - r_14_=−0.658, CI=−0.88 to −0.22, p=0.008). Therefore, even when self-ordered, there were substantial switching RT costs, the switching RT costs were optimised over time with learning, and individuals who made more switches had lower switch RT costs.

Repeat-trial RT varied between studies and with practise in a manner that indicated learning effects. A two-way rm-anova of repeat-trial RT showed main effects of study (F_1,29_=16.866, p<0.001, 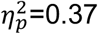), learning stage (F_1,29_=309.770, p<0.001, 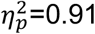) and a significant study * learning stage interaction (F_1,29_=14.516, p<0.001, 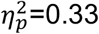). Repeat-trial RT was significantly slower in study 1 (two-sample t-test: t_29_=4.119, CI = 0.29 to 0.85, p<0.001, d=1.20) but decreased at a greater rate between the first and last learning stages (two-sample t-test: t_29_=−2.878, CI = −0.37 to −0.06, p=0.007, *d*=0.93).

### Task performance relates to structured behavioural strategies

Notably, the number of points scored did not relate in a trivial manner to the total switches made. This was also the case when controlling for the number of trials completed (see supplemental materials for more details). However, the algorithmic complexity of individuals’ behaviour (Figure 2c–d) had a significant negative correlation with points scored (partial Pearson: **Study 1** - r_15_=−0.646, CI=−0.87 to −0.22, p=0.017; **Study 2** - r_14_=−0.805, CI=−0.93 to −0.50, p=0.002) and the number of response errors made (partial Pearson: **Study 1** - r_15_=−0.695, CI=−0.89 to −0.31, p=0.004; **Study 2** - r_14_=−0.785, CI=−0.93 to −0.46, p<0.001). NB:-These partial correlations control for the number of events completed. That is, people whose behavioural routines were more structured and whose trial-responses were more accurate achieved higher scores. Exemplar behavioural timecourses from each study are rendered in Figure 2e–f.

As predicted, algorithmic complexity was significantly lower in study 2, compared to study 1 (two-sample t-test: t_29_=−4.500, CI=−0.36 to −0.14, p<0.001, *d*=1.26). Moreover, study 2 exhibited a significantly greater repetition bias than study 1 (two-sample t-test: t_29_=6.643, CI=0.19 to 0.36, p<0.001, *d*=1.53).

Furthermore, within-subject behaviour became increasingly structured as the SOS was practiced. Algorithmic complexity in the first behavioural learning stage negatively correlated with the sum of change across all three learning stages (partial Pearson **Study 1** - r_15_=−0.515, CI=−0.81 to −0.03, p=0.0496; **Study 2** - r_14_=−0.899, p<0.001, CI=−0.97 to −0.72). This indicates that increases in behavioural structure were greater for those who were initially behaving in a less structured manner.

Therefore, individuals developed more structured behaviour with practice, being characterised by ‘chunks’ of repeat-trials that were consistently punctuated by switches between rules and tasks, and behaviour in study 2 began at a more structured level, due to structured routines developed during the extensive pre-training.

### SOS activates multiple-demand cortex and reduces default mode network activation

We sought to identify brain regions where activation increased when individuals opted to switch between tasks or rules. Collapsing across switch-type, the cluster-corrected group-level t-contrast associated SOS with activation increases in the dorsolateral frontoparietal cortices, cerebellum, insula, putamen, and thalamus (Figure 3a). This pattern was consistent across studies (dice similarity = 0.58). The inverse t-contrast associated SOS with activation reductions in regions associated with the DMN (Figure 3b) and this pattern was consistent across studies (dice similarity = 0.56).

**Figure 3.**
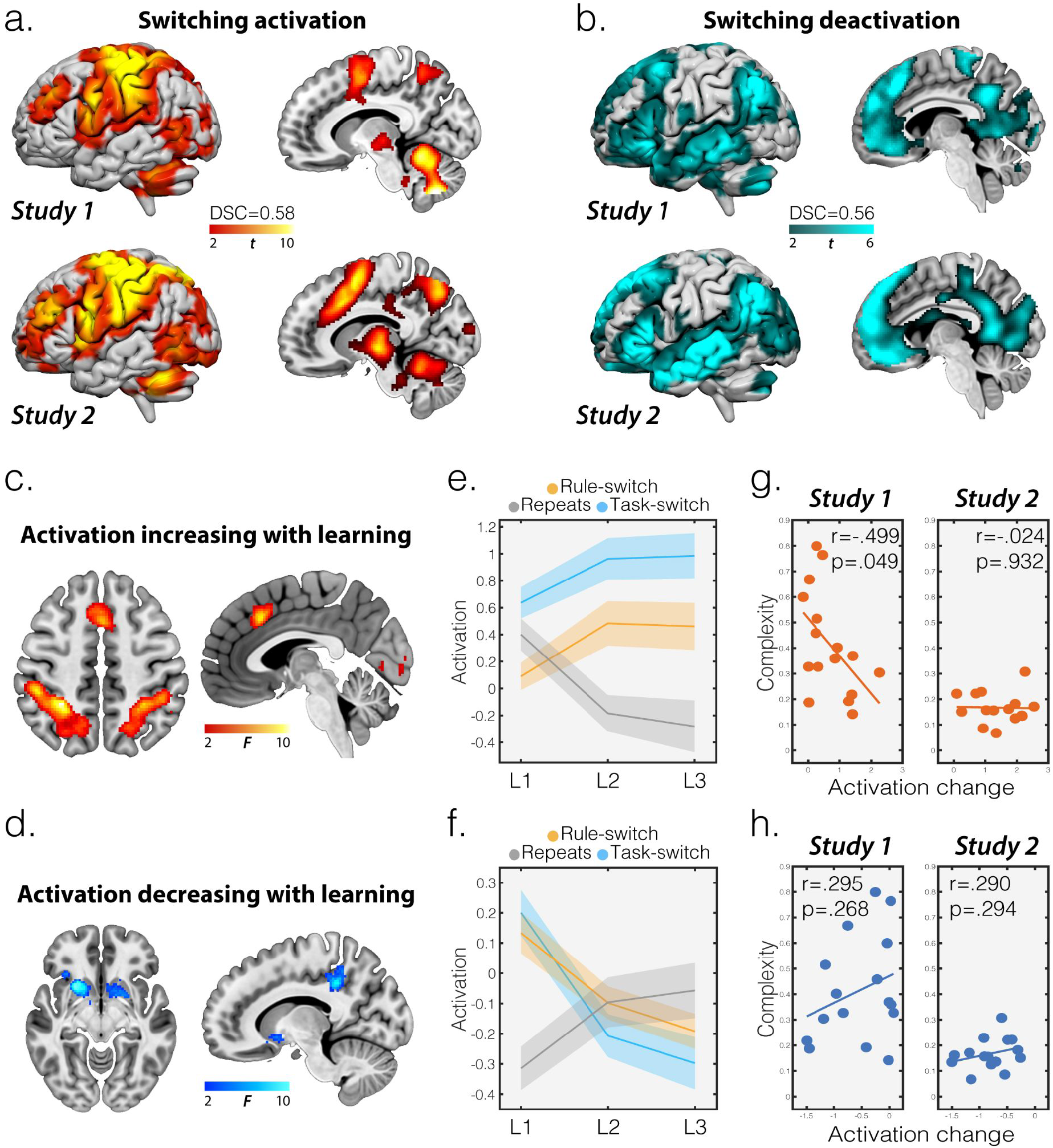
Multiple-demand and default mode activation varies with SOS learning. Group-level activation increases (a) and decreases (b) predicted by switching events. Dice similarity coefficients (DSC) represent the spatial similarity of significant voxels between the studies. d/e) Clusters of activation change over learning stages separated based on whether activation increased (c) or decreased (d). e/f) Mean activation independently examined for each learning stage (L) and condition for the MD (e) and DMN (f) composite ROIs (shaded area represents the standard error). g/h) Correlations between the sum of activation change over learning stage and normalised algorithmic complexity of SOS behaviour for the MD (g) and DMN (h) composite ROIs. All activation maps are thresholded at an uncorrected p<0.01, followed by a p<0.05 FDR cluster correction.

### Multiple-demand and default mode network activation changes with learning

Previous studies have examined activation changes when simple tasks are established through practise (Hampshire et al., 2016, 2019; Petersen et al., 1998; Ruge et al., 2019). To expand on this, voxelwise activation was compared across time during SOS. For each of the three learning stages, a 1st-level contrast was estimated for switching minus trial event activation. These contrasts were examined using group level random effects analyses with a 3 * 2 RM-ANOVA design where the learning stage was the within-subject factor and study was the between-subject factor.

The main effect of learning stage contrast rendered six bilateral regions where the difference between switching and trial activation significantly differed between the learning stages. For each cluster, a paired t-test determined the direction of the activation change using the mean bilateral activation from the 1st and last learning stages. Each cluster showed significant differences that survived Bonferroni correction (alpha=0.0083=0.05/6, see supplemental results). We combined those regions whose activation increased (Figure 3c) into a composite MD region of interest (ROI) of the anterior cingulate, dorsolateral parietal cortex and occipital cortex clusters. The MD ROI activation was significantly increased in the last learning stage, compared to the first (paired t-test: t_30_=7.230, CI=0.75 to 1.34, p<0.001, d=0.68). Those regions where activation decreased were combined into a composite DMN ROI and included the posterior cingulate, putamen and temporal lobe (Figure 3d). The DMN ROI activation was significantly decreased in the last learning stage, compared to the first (paired t-test: t_30_=7.432, CI=0.49 to 0.85, p<0.001, d=1.15).

Notably, MD activation increasing and DMN activation decreasing with learning is the exact opposite direction to what has been observed during simple stimulus-response learning tasks (Hampshire et al., 2019). To understand why this learning effect appeared to be inverted, we independently examined trial response activation and switching activation in the MD ROI across the learning stages (Figure 3e). The trial response events are more comparable to simple learning curve paradigms. Directly replicating these studies, activation in the MD ROI varied across the learning stages for trial responses (one-way RM-ANOVA: F_2,60_=16.043, p<0.001, 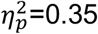) and was significantly decreased in the last learning stage, compared to the first (paired t-test: t_30_=−4.573, CI=−0.99 to −0.38, p<0.001, d=0.82). Conversely, MD ROI switching activation also varied across learning stage (one-way RM-ANOVA: **Task-switch** - F_2,60_=3.329, p=0.043, 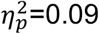; **Rule-Switch** - F_2,60_=6.313, p=0.003, 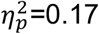) but was significantly increased in the last learning stage, compared to the first (paired t-test: **Task-switch** - t_30_=2.224, CI=0.03 to 0.67, p=0.034, d=0.40; **Rule-switch** - t_30_=2.669, CI=0.09 to 0.66, p=0.012, d=0.48).

The opposite pattern was evident for the DMN ROI activation (Figure 3f). Specifically, activation varied across the learning stages for trial responses (one-way RM-ANOVA: F_2,60_=10.119, p<0.001, 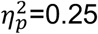) but was significantly increased in the last learning stage compared to the first (paired t-test: t_30_=3.595, CI=0.11 to 0.40, p=0.001). Conversely, DMN ROI switching activation also varied across learning stages (one-way RM-ANOVA: **Task-switch** - F_2,60_=16.544, p<0.001, 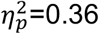; **Rule-switch** - F_2,60_=15.080, p<0.001, 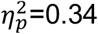) but was significantly decreased in the last learning stage, compared to the first (paired t-test: **Task-switch** - t_30_=−4.398, CI=−0.73 to −0.27, p<0.001, d=0.79; **Rule-switch** - t_30_=−5.112, CI=−0.46 to −0.20, p<0.001, d=0.92).

The sum of differences in mean MD activation over learning stage negatively correlated with individuals normalised algorithmic complexity (pearson, r_15_=−0.499, p=0.049, CI=−0.798 to - 0.004) in study 1 (Figure 3g) but not in study 2 (r_14_=−0.240, p=0.932, CI=−0.53 to 0.49). The equivalent analysis of the DMN ROI activation (Figure 3h) did not correlate with algorithmic complexity (pearson, **Study 1** - r_15_=0.295, p=0.268, CI=−0.24 to 0.69; **Study 2** - r_14_=0.290, p=0.294, CI=−0.26 to 0.70).

### Functional connectivity between frontoparietal networks and subcortical regions increases with learning

In accordance with the behavioural results showing that repeat-trial RT varied between studies and with practise (Figure 4a), the FC predicted by trial responses varied across learning stages in a manner that was largely specific to study 1. The mass-univariate glme modelling of the gPPI weighs across the learning stages identified connections whose FC significantly varied across learning stages. In study 1, the models that survived FDR correction for multiple comparisons were predominantly for connections between networks (Figure 4b) associated with cognition, such as the DMN, networks associated with MD (frontoparietal and attention networks), as well as with subcortical regions (Figure 4c). The glme models’ t-statistic was used to determine whether the FC increased or decreased across the learning stages. This indicated that FC increased as the SOS task was practised in study 1. Notably, these effects were largely absent in study 2 (Figure 4d).

**Figure 4.**
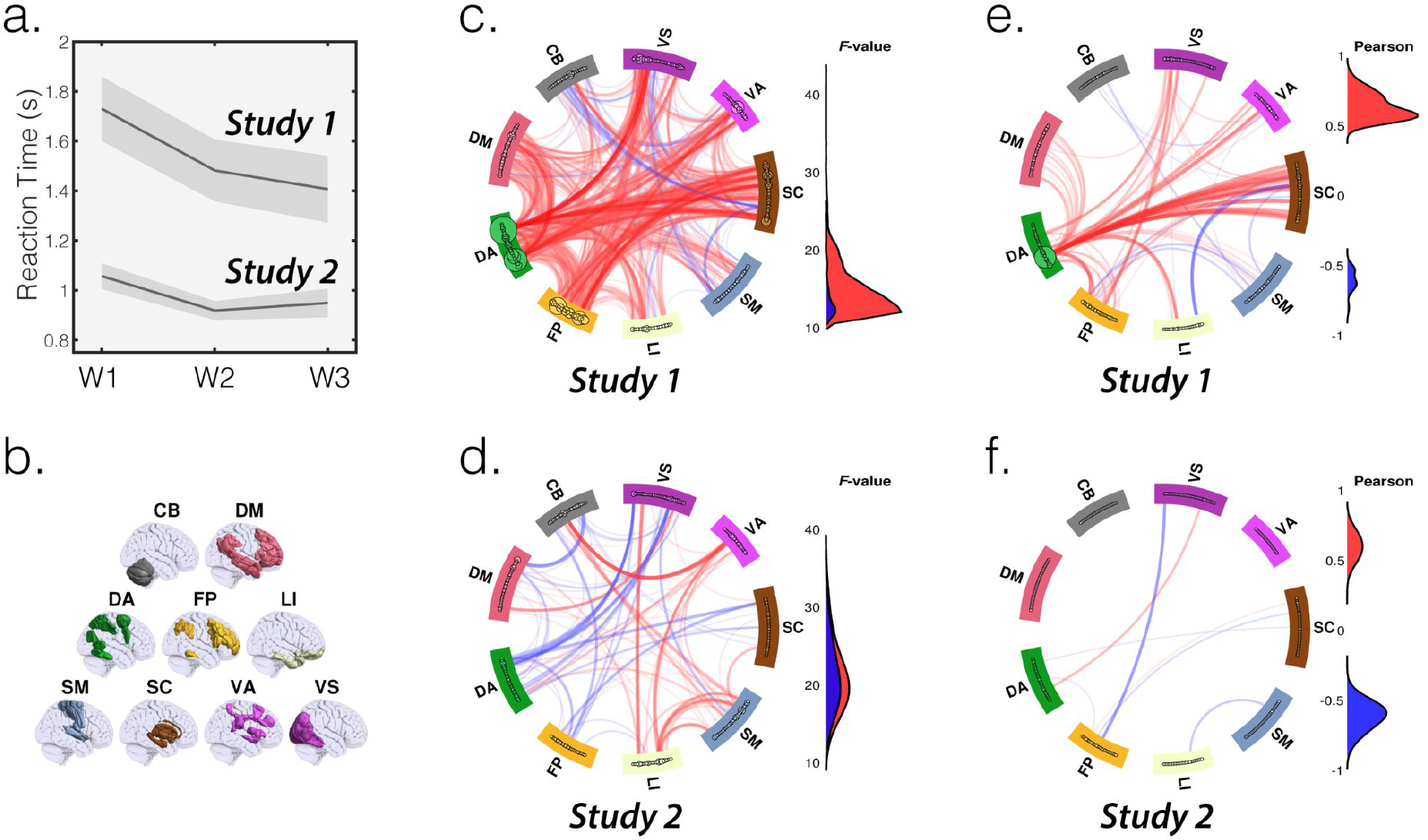
Increased between-network functional connectivity with learning. a) Reaction time (RT) decreased at a greater rate between the three windows (W) in study 1. The mean repeat-trial RT was plotted for each window with the standard error of the mean. b) Task-dependent functional connectivity (FC) changes for trial responses were estimated between brain regions and these were ordered by nine networks for visualisation purposes. c/d) Generalised linear mixed effects models identified connections whose strength significantly varied over the learning stages for study 1 and 2. The model t-statistic was used to render the connections in red or blue for positive or negative change, respectively. e/f) Partial correlations were used to estimate the relationship between changes in RT and FC over time for study 1 and 2 (controlling for the number of trials completed). Here, a positive correlation indicates an increase in response speed and FC. Each set of connectivities were FDR corrected and the surviving weights were rendered in the schemball and in the accompanying density distribution plot that was within-set normalised. CB=Cerebellum, DM=Default Mode, DA=Dorsal Attention, FP=Frontoparietal, LI=Limbic, SM=Somatomotor, SC=subcortical, VA=Ventral Attention, VS=Visual.

Finally, we estimated the relationship between the decreases in RT and FC over the learning stages. A mass univariate approach was taken where a partial correlation was calculated for each connection using the difference in gPPI and repeat-trial RT between the 1st and last learning stages. The partial correlation allowed this relationship to be estimated whilst controlling for the number of trials an individual completed. The resulting set of correlation coefficients were FDR-corrected for multiple comparisons. This unconstrained analysis showed significant correlations between the same systems identified in the previous analysis, demonstrating that a greater decrease in RT with learning correlated with a greater increase in FC. Again, these effects were specific to study 1 (Figure 4e) and largely absent in study 2 (Figure 4f).

## Discussion

The novel self-ordered switching paradigm developed here enabled new insights to be derived into the processes by which people learn complex multi-component tasks. In this executively challenging context, the learning process was characterised by the concurrent optimisation of behavioural and functional-network deployment; specifically, hierarchically structured behavioural routines that minimised switching costs were optimised based on the monitoring of multiple outcomes. Concurrently, functional-network resources were redeployed towards the executive switching points around which those routines were organised.

In everyday life we often are confronted with situations where multiple concurrent tasks must be performed. Furthermore, real-world tasks are often intrinsically hierarchical in their demands, requiring sub-goals to be completed in order to achieve overarching goals (Poljac et al., 2018). The majority of experimental paradigms that have been used to study cognitive control processes enforce careful control of the flow of events, for example, specifying when the participant must switch between sub-tasks (Kiesel et al., 2010; Koch et al., 2018). These types of design facilitate the careful balancing of conditions, but they lack ecological validity insofar as they do not provide the participant with the opportunity to apply their executive abilities in order to organise their own behaviour (Goldberg & Podell, 2000). This is important, because structured routines are a defining characteristic of human behaviour with disorganisation being symptomatic across prominent neurological and psychiatric populations (Averbeck et al., 2011; Collins & Koechlin, 2012; Fallon et al., 2013).

Moreover, the majority of cognitive control studies do not take into account the shifting role that brain systems play as a task is learnt. We argue that this is a critical oversight because the purpose of cognitive control systems is not just to support complex behaviour, but more to enable the learning of such behaviours in order that they can be performed with minimal effort.

Indeed, we and others have previously reported that the changes in brain activity and connectivity that are associated with learning during commonly applied cognitive control paradigms can dwarf the control processes that those paradigms were designed to examine (Badre et al., 2010; Erika-Florence et al., 2014). More specifically, in a recent sequence of articles, we and others reported that the learning of simple stimulus-response mappings from either instruction or feedback was accompanied by an ongoing reduction in event-related activity within frontoparietal networks associated with effortful cognition and a reduction in default mode network activity (Hampshire et al., 2016, 2019; Ruge et al., 2018, 2019; Ruge & Wolfensteller, 2010). This likely reflected a steady transition from deliberate to automated modes of behaviour.

The results of the current study extended that work by examining the shifting role of brain networks during the learning of tasks that are more ecologically valid insofar as they require multiple sub-tasks to be efficiently self-organised. When this learning process was supported by detailed ongoing feedback in study 1, the behavioural routines that participants tended to develop had several notable characteristics, these being the minimisation of number of switches performed, the hierarchical sequencing of task and rule switches, and the tendency towards algorithmically simpler sequences with practice. Interestingly, all three of these characteristics correlated with overall task performance, as quantified in number of points awarded across learning stages or in terms of individual differences in behavioural routines; however, from a simple computational perspective this would not necessarily have been the case.

For example, task switches could have been performed at the same frequency, or subordinate to rule switches whilst producing the same number of points per trial completed. However, participants in study 1 tended to organise the rule switches subordinate to the task switches, a structure that was implicitly encouraged by the structure of the feedback elements. Notably, task switches were more effortful, as evidenced by greater response time costs; therefore, the bias towards more rule than task switches, and fewer switches overall, was adaptive insofar as it reduced overall executive costs. Relatedly, the application of algorithmically simpler routines was demonstrably less effortful, being accompanied by reduced switching costs and the achievement of higher overall scores. From this perspective, the development of hierarchically structured behaviour can be seen to reflect a tendency to optimise towards a mode of behaviour that requires minimal switching effort to produce maximum reward.

Indeed, evidence of a tendency to self-order behaviour in a structured manner even when such structure is not a task requirement has been seen previously in some previous studies. A prominent example is voluntary task-switching (VTS) paradigms, which instruct participants to randomly choose between two tasks on each trial. Interestingly, the original VTS reports imply that participants did not fully adhere to this instruction and exhibited a significant repetition bias (Arrington & Logan, 2004, 2005). This is evidence of the behavioural self-ordering that reduced the frequency of task-switching that may have resulted in an advantageous reduction in cognitive demand. Similarly, here, participants who were better practised at SOS (study 2) exhibited a significantly greater trial repetition bias, and also behaved with a more structured routine.

Relatedly, we have also observed that when identifying targets during a novel self-ordered intradimensional-extradimensional switching paradigm, participants tend to hierarchically order their search behaviour between stimuli of different dimensions, this in a context where there is no intrinsic computational advantage to sequencing the search by stimulus dimension but switching between dimensions nonetheless incurs a greater response time costs (Hampshire & Owen, 2006).

Perhaps the most striking result of the current study pertains to how the underlying network resources associated with repeat and switch trials changed as the behavioural routine was established. More specifically, the optimisation of a structured behavioural routine was concomitant with a progressive ‘fine-tuning’ of the brain’s functional activation that was more complex in its dynamics than that which has previously been observed during simple stimulus-response learning (Hampshire et al., 2016, 2019; Ruge et al., 2019; Ruge & Wolfensteller, 2010). In accordance with those past findings, frontoparietal activation in response to the trials was observed to decrease as they were practised whilst the degree of default mode network deactivation was also seen to reduce. Unexpectedly though, we also observed the opposite pattern of change for switching events, with frontoparietal activation increasing and DMN activation decreasing as a function of learning. These executive switching events were effortful and critical for guiding behaviour and had to be conducted in a strategic manner. This distinguishes the SOS from most other task-switching and VTS paradigms.

Furthermore, the ACC and DLPFC regions identified here are part of the MD system and generally activate in response to increasing cognitive effort (Crittenden & Duncan, 2014). Therefore, this pattern of results likely reflects a redeployment of flexible and limited capacity network resources towards the executive switching events that enable structured behaviour as the simple stimulus-response mappings that constitute individual trials become automatic through practice. One intriguing possibility is that frontoparietal systems operate at the leading edge of the automatisation process, enabling progressively more complex hierarchical behaviours to be developed. A testable prediction of this is that the switching activation would also decrease with enough practise. This would make an interesting prediction that could be tested across multiple SOS scanning sessions, perhaps with paradigms that have scope for deeper hierarchies that limited capacity resources can progressively develop.

In terms of individual differences, it is interesting to note that in study 1, the scale of change in trial vs switch activation over learning stages related to the complexity of the behavioural routine. Those participants who could establish an optimal balance between repeating a condition and making the costly switches behaved in a more structured manner, scored the most points and exhibited a greater ‘fine-tuning’ of frontoparietal activation away from simple trials to the executive switching events. This indicates that optimal structured routines may also help accelerate the learning process.

The key difference between study 1 and study 2 was that the latter had more substantial training prior to entering the scanner and only minimal feedback within the scanner. These modifications were intended to provide a control condition where the potential for further optimisation of the behavioural routine was greatly reduced, with participants in study 2 being likely further along the ‘learning curve’. The behavioural results supported this view, as in study 2, SOS was performed faster and with routines that had low algorithmic complexity. In this context, the changes in behaviour across time also were both less pronounced. RT dropped at a lower rate and the algorithmic complexity progressed towards a more stereotyped pattern that expressed more consistently repeating structures.

Intriguingly, the differences across studies in behavioural and network activity signatures of learning was also evident in the connectivity analyses. More specifically, in study 1 we observed widespread increases in FC between functional networks in the brain that have been associated with cognition and their connections with subcortical regions. A similar increase in network coherence was evident in the individual differences analysis, where greater reductions in RT were associated with a greater increase in network functional connectivity with learning. In contrast, FC changes with practise were largely absent in study 2. These findings accord with the notion that cognitive behaviours can through practice be supported by higher efficiency network states that are characterised by lower activity but increased connectivity (Hampshire et al., 2016; Soreq et al., 2019, 2021; Stokes, 2015).

More broadly, we believe that paradigms such as the SOS that enable participants the freedom to self-organise their behaviour have untapped potential for understanding normal and pathological cognitive control functions. The natural variation in the behavioural efficiency of healthy adults suggests that there is a potential to examine self-ordered behaviours in patient populations that suffer from cognitive impairments. Indeed, high activity-low connectivity states have been reported in some dysexcutive populations when perfoming tasks that require cognitive control (Hampshire et al., 2013; Sheffield et al., 2021). Furthermore, it could be predicted that patients may take longer to generate a stable behavioural routine and that this routine would be more complex, inefficient and refined at a slower rate, compared to healthy controls. These features may dissociate across patient populations for example, Parkinson’s patients show difficulty in updating strategies during set-shifting and can show a reduced tendency towards structured dimensional switching routines (Fallon et al., 2013), whereas patients with schizophrenia may prematurely implement a strategy before sufficiently exploring the options (Averbeck et al., 2011). Future work also should focus on the learning of structured routines over longer time frames and within paradigms that provide even greater potential for hierarchical self-organisation and should examine how the responsiveness of functional networks evolves as deeper hierarchical behaviours are established.

## Acknowledgements

Richard E. Daws is supported by an EPSRC MRes/PhD scholarship at the Imperial College Centre for Neurotechnology (EP/L016737/1).

## Data/code availability statement

The MRI, behavioural data and the analysis and visualization codes will be made available via online repositories upon acceptance for publication. The SOS paradigm task codes are available upon request.

## CRediT author statement

**Richard E. Daws:** Conceptualisation, Formal analysis, Methodology, Visualisation, Writing - Original Draft, Review & Editing. **Gregory Scott:** Conceptualisation, Software, Writing - Original Draft, Review & Editing. **Eyal Soreq:** Conceptualisation, Visualisation. **Robert Leech:** Conceptualisation, Supervision, Writing - Original Draft, Review & Editing. **Peter J. Hellyer:** Conceptualisation, Writing - Original Draft, Review & Editing. **Adam Hampshire:** Conceptualisation, Software, Supervision, Project administration, Funding acquisition, Writing - Original Draft, Review & Editing.

## Footnotes

1 “SPM12 - Wellcome Trust Centre for Neuroimaging - UCL.” Accessed July 30, 2019. https://www.fil.ion.ucl.ac.uk/spm/software/spm12/.

2 “MATLAB - MathWorks - MATLAB & Simulink.” Accessed July 30, 2019. https://www.mathworks.com/products/matlab.html.

